## Supplementary material for "Brain microRNAs among social and solitary bees": Table S1, Table S2, Table S3, Metadata for Tables S4-S8

**Table S1.** Genome assemblies used for alignment of small RNA sequences or genome scans for known small RNAs. The last six species were used only for genome scans.

| Species | Genome assembly | Reference |
|---|---|---|
| *Apis mellifera* | Amel v4.5 | Elsik et al. 2014 |
| *Bombus impatiens* | Bimp v2.0 | Sadd et al. 2015 |
| *Bombus terrestris* | Bter v1.0 | |
| *Megalopta genalis* | Mgen v1.0 | Kapheim et al., unpublished (NCBI PRJNA494872) |
| *Megachile rotundata* | Mrot v1.0 | Kapheim et al. 2015 |
| *Nomia melanderi* | Nmel v1.0 | Kapheim et al. 2019 |
| *Apis florea* | Aflo v1.0 | Baylor College of Medicine, unpublished (NCBI PRJNA45871) |
| *Dufourea novaeangliae* | Dnov v1.0 | Kapheim et al. 2015 |
| *Eufriesea mexicana* | Emex v1.0 | Kapheim et al. 2015 |
| *Habropoda laboriosa* | Hlab v1.0 | Kapheim et al. 2015 |
| *Lasioglossum albipes* | Lalbi v2 | Kocher et al. 2013 |
| *Melipona quadrifasciata* | Mqua v1.0 | Kapheim et al. 2015 |

**Table S2.** Sample acquisition and library preparation methods. Samples were collected into liquid nitrogen and stored at -80 °C until dissection. All libraries were prepared from RNA isolation from whole brain tissue. All libraries were sequenced for 51 cycles on a HiSeq 2500. *B. impatiens*, *M. genalis*, *N. melanderi* libraries were pooled and sequenced in one lane. *A. mellifera*, *B. terrestris*, and *M. rotundata* libraries were pooled and sequenced in one lane.

| Species | Collection | Sample type | RNA Isolation | RNA Quality Assessment | Sequencing Center | Library Prep | # reads | Reads mapped to genome |
|---|---|---|---|---|---|---|---|---|
| *Bombus impatiens* | Commercial colony (BioBest, Romulus, MI, USA) | Worker | mirVana miRNA Isolation kit with phenol (Ambion) | TapeStation (Agilent) – USU Center for Integrated Biosystems | University of Illinois Roy J. Carver Biotech Center | Illumina TruSeq Small RNA Sample Preparation kit | 18,173,120 | 11,147,865 (61.3%) |
| *Megalopta genalis* | Barro Colorado Island, Panama[¥] | Lab-reared female | | | | | 17,273,381 | 14,459,264 (83.7%) |
| *Nomia melanderi* | Touchet, WA, USA | Reproductive female | | | | | 21,916,316 | 16,681,729 (76.1%) |
| *Apis mellifera* | Urbana-Champaign, IL; Tyson Research Station, St. Louis, MO, USA | Worker | TRIzol reagent (Thermo Fisher Scientific) | Bioanalyzer (Agilent) - Washington University Genome Technologies Access Center | Washington University Genome Tech Access Center | Illumina TruSeq, Clontech SMARTer small RNA library kit | 12,793,471 | 7,638,748 (59.7%) |
| *Bombus terrestris* | Commercial colony (Pollination Services Yad-Mordechai, Kibbutz Yad-Mordechai, Israel) | Worker | | | | | 16,270,644 | 11,205,739 (68.9%) |
| *Megachile rotundata* | Logan, UT, USA | Reproductive female | | | | | 19,160,796 | 15,357,334 (80.1%) |

[¥]*M. genalis* samples were exported under permit SEX/A-37-15

**Table S3.** Known microRNAs used in miRDeep2 microRNA detection protocol.

| Species | Source | Reference |
|---|---|---|
| *Apis mellifera* | miRBase v21 | Kozomara and Griffiths-Jones 2014 |
| *Drosophila melanogaster* | | |
| *Nasonia vitripennis* | | |
| *Tribolium castenum* | | |
| *Bombyx mori* | | |
| *Apis mellifera* | Small RNA sequencing | Ashby et al. 2016 (Table S1) |

**Table S4.** Gene models used for localization of microRNAs and predicted target analysis.

| Species | Genome annotation | Reference |
|---|---|---|
| *Apis mellifera* | Amel OGS v3.2 | Elsik et al. 2014 |
| *Bombus impatiens* | Bimp OGS v1.0 | Elsik et al. 2016 |
| *Bombus terrestris* | Bter v1.3 | Sadd et al. 2015 |
| *Megalopta genalis* | Mgen v1.0 | Kapheim et al., unpublished |
| *Megachile rotundata* | Mrot v1.1 | Kapheim et al. 2015 |
| *Nomia melanderi* | Nmel v1.0 | Kapheim et al. 2019 |

**Table S5 (separate file)**. Enrichment results for predicted targets of lineage-specific miRs and "social genes". Includes gene lists, conversion lists based on reciprocal blastp results, and overlap test statistics. Column head descriptions are in 'ColumnDetails' sheet.

**Table S6 (separate file).** Results from genome scans for miRNA seed matches in Rfam. Descriptions for each sheet and column header are provided in the 'Metadata' sheet.

**Table S7 (separate file).** Final miRNA sets for each species. Descriptions for each sheet and column header are provided in the 'Metadata' sheet.

**Table S8 (separate file).** Predicted targets and orthogroup ages of lineage-specific microRNAs in each species. Descriptions for each sheet and column header are provided in the 'Metadata' sheet.

**Figure S1 (separate file).** Predicted targets of lineage-specific miRNAs in relation to social behavior. Genes that are both predicted targets of lineage-specific miRNAs and genes with differential expression in a social context (solid outlines) or genes under selection (dashed outlines) are represented by overlapping circles for each study and species. Numbers of lineage-specific miRNA targets are given for each species. Colors indicate different studies. Overlaps not significantly different from random (representation factor, RF=1) are unlabeled, while significant over- or under-enrichments are marked with asterisks with RF and p-value as indicated.
