## Supplementary figures and images for "Brain microRNAs among social and solitary bees"

### Fig. S1

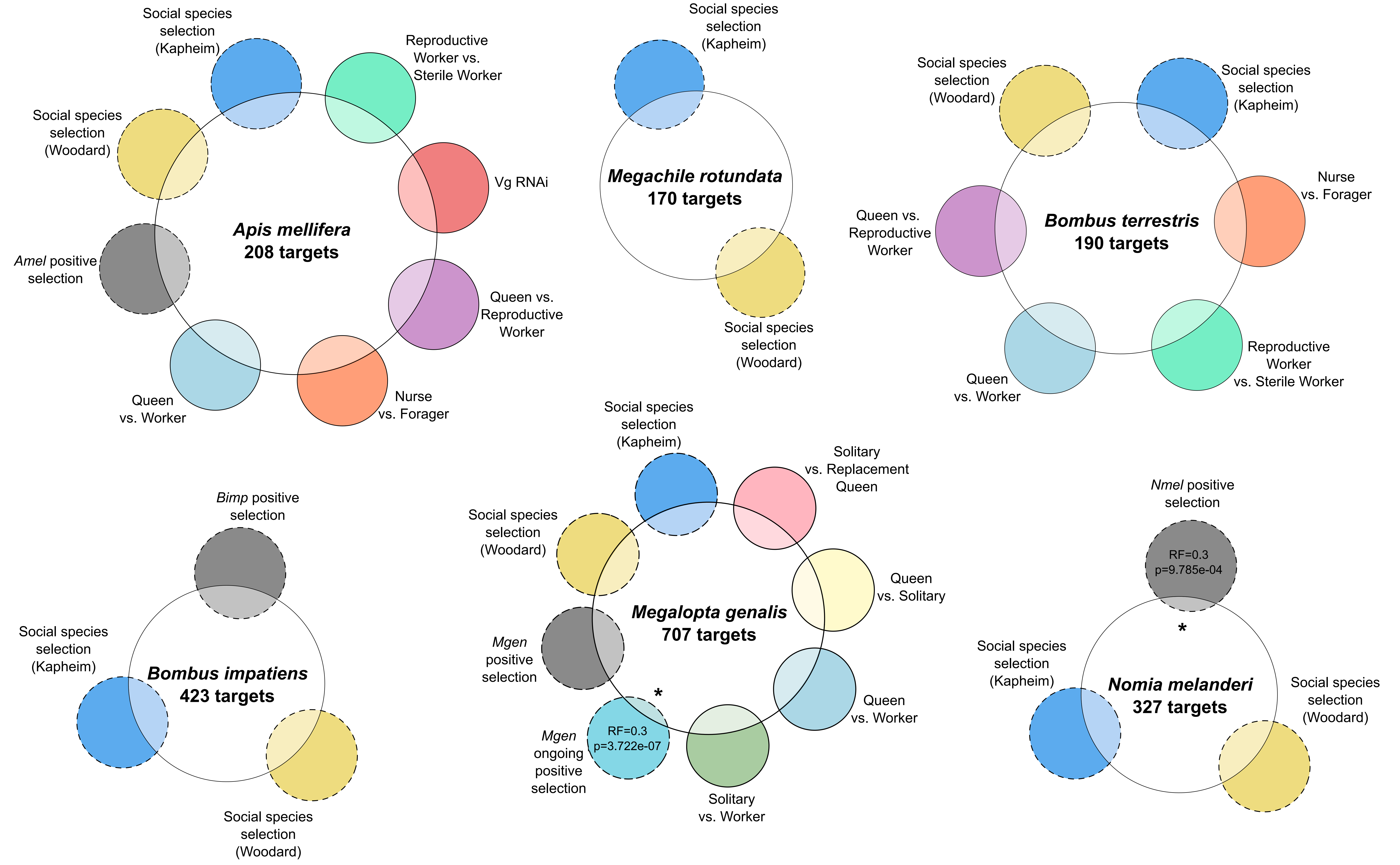
